# Omega-3 polyunsaturated fatty acids modify glucose metabolism in THP-1 monocytes

**DOI:** 10.1101/2024.03.01.582966

**Authors:** Michael J. Byun, Roni Armon, Tamaris Souza, Hope D. Anderson, Ayesha Saleem, Samantha D. Pauls

## Abstract

Chronic inflammation is a driving factor in diseases like obesity and type 2 diabetes. Enhanced glucose metabolism, including *via* oxidative phosphorylation, may contribute to heightened immune activation. A recent clinical trial showed that supplementation with the n-3 fatty acid α-linolenic acid (ALA) reduced oxidative phosphorylation rates in human monocytes. However, the mechanism remains unknown. Therefore, our objective was to explore the direct effects of ALA and docosahexaenoic acid (DHA) on glucose metabolism in a cell culture model and to explore the molecular mechanism.

THP-1 monocytes were treated for 48h with 10-40 μM of ALA or DHA and compared with vehicle and oleic acid controls. The Seahorse XFe24 system was used to approximate catabolic rates in the presence of glucose, including glycolysis and oxidative phosphorylation. The latter was validated by respirometry using an Oroboros O2k Oxygraph. Both ALA and DHA treatments reduced oxidative phosphorylation and increased glycolytic rates relative to control conditions. We identified pyruvate dehydrogenase kinase 4 (PDK4), an enzyme that inhibits the conversion of pyruvate to acetyl-CoA, as a possible mechanistic candidate. This gene was significantly upregulated by ALA and, to a greater extent, by DHA. Using fluorescent indicators, we also found that DHA increased reactive oxygen species while ALA had no effect.

Our data suggest that ALA and DHA trigger a re-wiring of bioenergetic pathways in monocytes, possibly *via* the upregulation of PKD4. Given the close relationship between cell metabolism and immune cell activation, this may represent a novel mechanism by which n-3 fatty acids modulate immune function and inflammation.

## Introduction

Obesity is a global epidemic with high prevalence not only in adults but also in children (Han and Lean, 2016; Rao et al., 2016). In Canada, approximately 30% of children aged 5-17 were classified as overweight or obese (Rao et al., 2016). This represents a significant risk in the early development of adverse metabolic characteristics which promote the initiation and progression of chronic metabolic diseases such as type 2 diabetes mellitus (T2DM) and cardiovascular disease (CVD) (Han and Lean, 2016; Park et al., 2023). One of these characteristics is chronic inflammation (Natto et al., 2019; Hildebrandt et al., 2023). Inflammation is an essential response that is initially triggered and regulated by innate immune cells such as monocytes during an infection or injury. However, its chronicity derives from an improper resolution of the inflammatory response (Daryabor et al., 2020). Therefore, early intervention strategies that can slow the progression of obesity-associated metabolic diseases hastened by inflammation could be vital in improving long-term health outcomes and quality of life for affected patients.

Monocytes are the most common innate immune cell type in the blood. They contribute to a chronic inflammatory phenotype by secreting cytokines directly and by migrating to sites of infection, damage, or dysfunction (eg. hypertrophic adipose tissue) where they differentiate into macrophages. (Hildebrandt et al., 2023; Daryabor et al., 2020). Recent studies have shown that chronic inflammation is associated with a dysfunction in monocyte bioenergetics or energy-generating pathways (Hartman et al., 2014; Pauls et al., 2021). For example, oxidative phosphorylation, the process by which pyruvate is catabolized in the mitochondria *via* the Kreb’s cycle, coupled with the electron transport chain (ETC) for ATP production, is heightened in peripheral blood mononuclear cells (PBMC) from patients with T2DM (Hartman et al., 2014), and is correlated to severity of obesity (Pauls et al., 2021). Targeting this mitochondrial dysfunction may alter the course of inflammation-associated metabolic disorders, for better or for worse.

Omega-3 polyunsaturated fatty acids (n-3 PUFAs) such as plant-derived alpha-linolenic acid (ALA) and fish oil-derived docosahexaenoic acid (DHA) and eicosapentaenoic acid (EPA), are natural products with high therapeutic potential. A recent meta-analysis reported that higher n-3 PUFA circulating levels is linked to overall lower risk of mortality (Harris et al., 2021). This protective effect is partly attributed to the dampening of pro-inflammatory markers (Baranowski et al., 2012; Browning et al., 2007; Faintuch et al., 2007; Itariu et al., 2012); however, the differences between ALA, DHA and EPA and their mechanisms of action are not well-understood.

A recent clinical trial demonstrated that 4g/day of ALA but not DHA supplementation for 28 days in women with obesity reduced oxidative glucose metabolism in monocytes, measured by oxygen consumption rate (OCR) (Pauls et al., 2021). Here, we sought to test the direct effects of ALA and DHA on glucose metabolism in THP-1 monocytes and to identify candidate genes that may mediate the effects.

## Methods

### Cell culture and n-3 PUFA treatment

THP-1 human monocytes (TIB-202, American Type Culture Collection) were maintained between 2-10×10^5^ cells/ml in Roswell Park Memorial Institute 1640 (RPMI-1640, Thermo Fisher Scientific) medium containing 10% FBS (Gibco) and 0.1% penicillin-streptomycin (Gibco). Cells were cultured at 37°C and 5% CO2 in a humidified incubator. All fatty acid stock solutions, including ALA, DHA, and oleic acid (OA), a monounsaturated fatty acid control (Cayman Chemicals) were adjusted to 100 μM in ethanol and flushed with nitrogen to prevent oxidation. Prior to cell treatments, fatty acids were conjugated to 1% fatty acid-free bovine serum albumin (BSA, MP Biomedicals) in PBS (Millipore Sigma) or, for later experiments, in pre-made fatty acid-free, low endotoxin 1% BSA solution in PBS (Millipore Sigma) for 1 h in a 37°C water bath. Cells were treated with vehicle control (ethanol) or 10-40 μM of fatty acid, as indicated, for 48h with an additional bolus dose at 24h.

### RT-qPCR

THP-1 monocytes were harvested and lysed for RNA extraction using the Monarch Total RNA Miniprep kit (New England Biolabs), according to the manufacturer protocol. cDNA was synthesized *via* reverse transcription from ~500 ng of RNA using LunaScript RT Supermix (New England Biolabs). Transcripts of interest were subsequently quantified using Luna qPCR master mix (New England Biolabs) and a Quant Studio 6 Flex instrument, following the 2^-ddCt^ method with β-actin as the reference gene (Maeß et al., 2010). Primer sequences (Integrated DNA Technologies) are shown in Table 1. Amplicon size was verified *via* agarose gel electrophoresis at least once for each target.

**Table 1.**
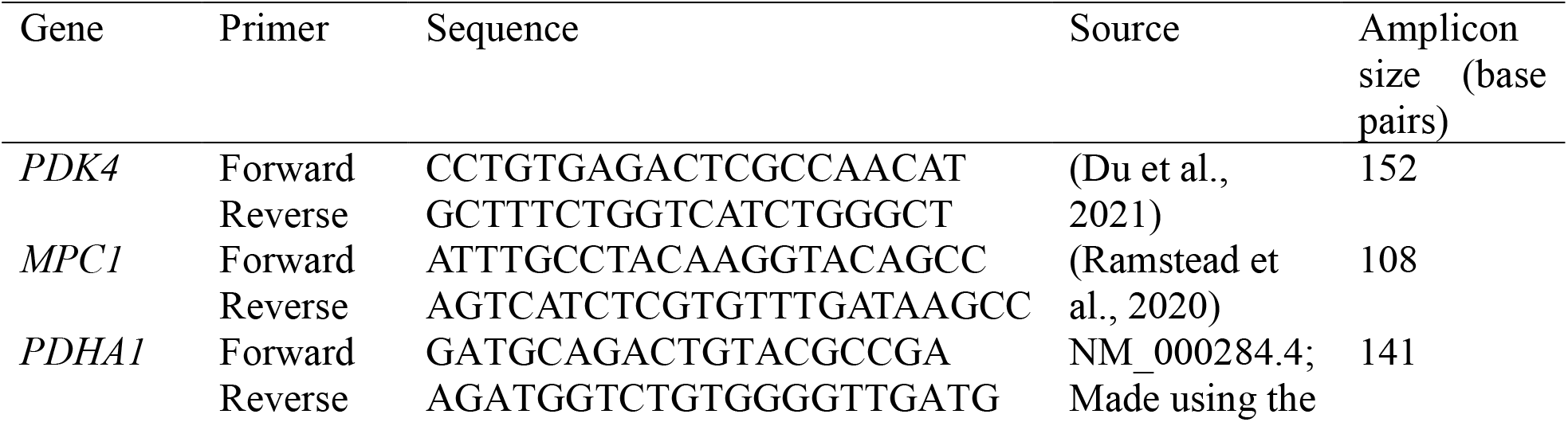
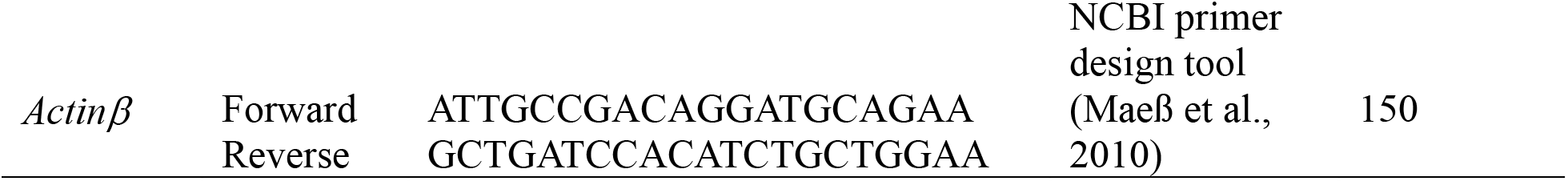
RT-qPCR primers.

### Seahorse XF analysis

XFe24 culture plates were coated with 35 μg/ml of CellTak adhesive (Corning) in 0.1 M of sodium bicarbonate (pH 7.4) for 30 min at room temperature. Plates were then washed twice with sterile deionized water. THP-1 monocytes were counted with a hemocytometer, centrifuged, and resuspended in complete Seahorse XF medium [Seahorse XF RPMI medium (Agilent Technologies) supplemented with 2 mM of L-Glutamine (Hyclone), 1 mM of sodium pyruvate (Gibco), and 10 mM of glucose (Gibco)]. Cells were plated at 2×10^5^ cells/well and the plate was centrifuged at 300 g for 5 minutes at room temperature to promote adhesion before incubating for 30 minutes at 37°C. Then the medium was replaced with 500 μL of fresh complete Seahorse XF medium. The plate was incubated for an additional hour in a CO2-free incubator. Mitochondrial stress test or ATP rate assay protocols were executed using a Seahorse XFe24 analyzer (Agilent Technologies), which includes the injection of electron transport chain inhibitors. These were 1.0 μM of the ATP-synthase inhibitor oligomycin (Millipore Sigma), 1.5 μM of the uncoupler carbonyl cyanide 4-(trifluoromethoxy) phenylhydrazone (FCCP; Millipore Sigma), and a mixture of 1 μM of the complex III inhibitor antimycin A + 0.1 μM of the complex I inhibitor rotenone (Millipore Sigma)]. Validation experiments were performed using N5,N6-bis(2-Fluorophenyl)[1,2,5]oxadiazolo[3,4-b]pyrazine-5,6-diamine (BAM15, Millipore Sigma) as an alternative uncoupling agent. Bioenergetic parameters such as oxygen consumption rates (OCR) and extracellular acidification rates (ECAR) were calculated using WAVE software (Agilent Technologies) and visualized using GraphPad Prism.

### Oroboros Oxygraph respirometry analysis

THP-1 monocytes were counted and loaded into an Oroboros O2k Oxygraph (Oroboros Instruments) at 2.5×10^6^ cells per chamber in 500 μL of fresh serum-free RPMI-1640 (Gibco, contains 11mM glucose and 2mM L-glutamine). After air calibration, the instrument’s Clarke-type oxygen electrodes were used to measure respiration rates in cells at 37 °C (Roy Chowdhury et al., 2020). O2 concentration and its background-corrected negative time derivative (ie. OCR) were monitored and recorded in real-time at baseline and after manual injection with 2μM oligomycin, 1μM of FCCP (2 or more injections), and 2μM antimycin A. Oroboros DatLab software was used to calculate the OCR parameters and the data were replotted using GraphPad Prism for visualization.

### Flow cytometry

MitoSOX Red (Thermo Fisher Scientific) and CellROX Green (Thermo Fisher Scientific) dyes were prepared and administered to cells according to manufacturer protocols, in separate experiments and fluorescence was quantified by flow cytometry (CytoFLEX LX). For MitoSOX red, cells were adjusted to 1×10^6^ cells/tube for staining with 5 μM dye; fluorescence acquisition was in the R-phychoerythrin (PE) channel. For CellROX Green, cells were adjusted to 5×10^5^ cells/tube for staining with 500nM dye; fluorescence acquisition was in the fluorescein isothiocyanate (FITC) channel.

### Statistical analysis

Analysis and graph plotting were performed using Prism software (version 10.0.3, GraphPad Software Inc.). Numerical data are expressed as mean±standard error of the mean (SEM). Statistical analyses included unpaired t-test for Oroboros experiments and one-way ANOVA followed by Tukey’s post-hoc test for all other experiments. Differences were considered significant at p<0.05.

## Results

### Direct treatment with ALA results in lower oxidative phosphorylation rates in the presence of glucose

In the clinical trial reporting a dampening of oxidative metabolism by ALA in monocytes, the n-3 PUFA was administered orally (Pauls et al., 2021). PUFA supplementation increases the levels of that PUFA in nearly every organ and tissue (Azab et al., 2020; Leng et al., 2017), including the circulating non-esterified (free fatty acid) form. However, many other systemic changes are expected, therefore we sought to understand whether the effect of ALA on monocyte metabolism was direct.

We first performed an ALA-dose curve (10 – 40 μM) experiment to identify the optimal concentration for treatment. We used the Seahorse X24Fe platform to perform a mitochondrial stress test, which measured bioenergetic parameters such as basal respiration, ATP-linked oxygen consumption rate (OCR) and extracellular acidification rate (ECAR). Qualitatively, we found concentration-dependent changes in all 4 bioenergetic parameters. A dose of 40 μM ALA reached significance compared to control (0 μM): Basal respiration had decreased by ~40%, ATP-linked OCR had lowered by ~60%, and ECAR had increased by ~60%, (**Figure 1**).

**Figure 1.**
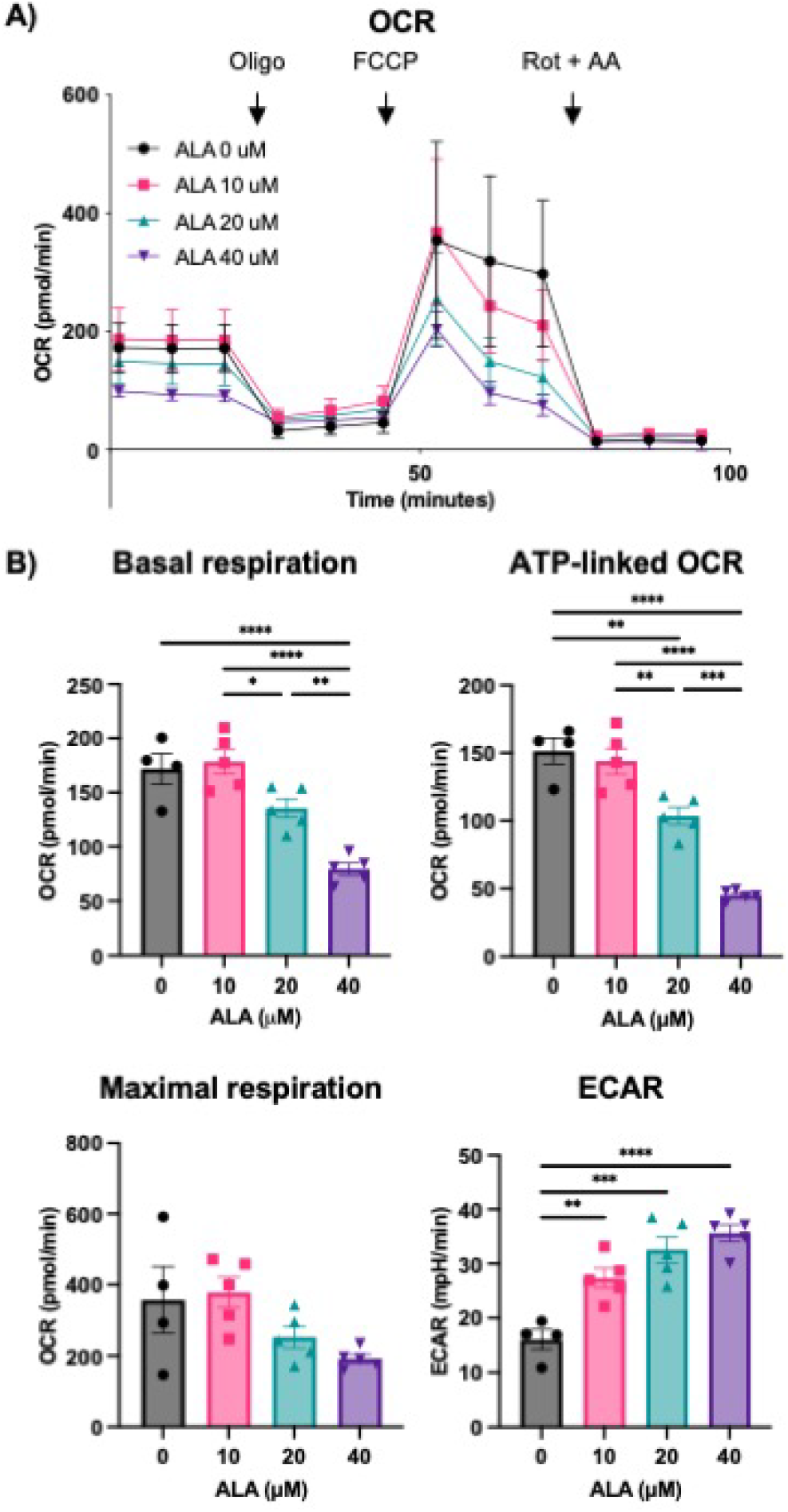
ALA alters bioenergetic parameters of THP-1 cells in a dose-dependent manner. Human THP-1 cells were treated with increasing concentrations of ALA (0 μM, 10 μM, 20 μM, 40 μM) in triplicates for 48 h with a bolus dost at 24 h. Bioenergetics were assessed by a Seahorse XF Mito Stress Test, performed on a Seahorse XFe24 analyzer. (A) Time plot showing changes in Oxygen consumption rate (OCR) following the injection of Oligomycin (oligo, 1 μM), uncoupling agent (FCCP, 1.5 μM) and Rotenone + Antimycin A (Rot, 0.1 μM + AA, 1 μM). Glucose was used as the metabolic substrate at a concentration of 10 mM. (B) Comparison of Basal respiration, ATP-linked OCR, Maximal respiration and Extracellular acidification rate (ECAR) between treatments. Data are shown as mean ± SEM of n=4-6 technical replicate wells and analyzed via One-Way ANOVA; * P < 0.05, ** P < 0.01, *** P < 0.001, **** P < 0.0001. Results are representative of two independent experiments.

We sought to validate this finding using the gold standard method – direct O2 measurement by respirometry using an Oroboros O2k Oxygraph instrument. Cells treated with vehicle or 40 μM ALA for 48 h, as above, were loaded into the instrument chambers (alternating which treatment was loaded into chamber A or chamber B) to follow oxygen consumption rates. Similar to the Seahorse XFe results, ALA treatment resulted in reduced ATP-linked respiration and maximal respiration rates (**Figure 2**) compared to vehicle control.

**Figure 2.**
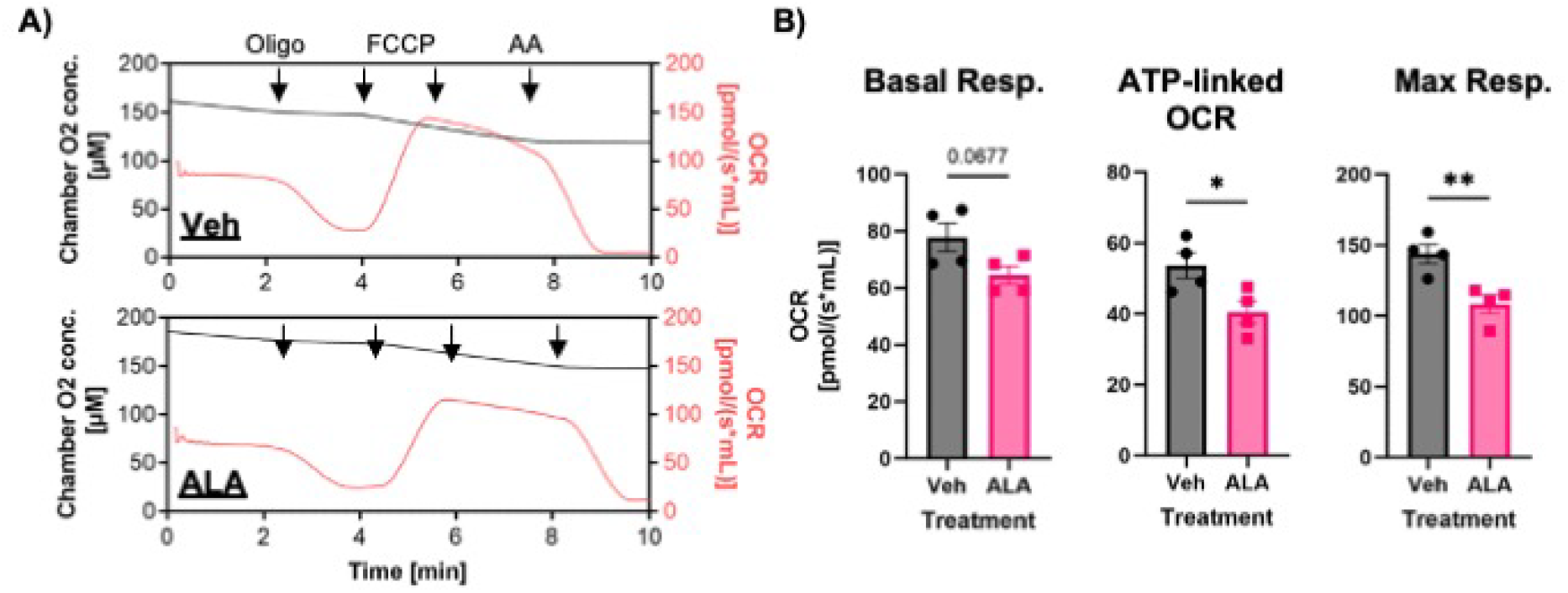
ALA reduces the oxygen consumption rate in monocytes as measured by high-resolution respirometry. THP-1 cells were treated with Vehicle control or 40 μM ALA as in Figure 1, then assessed for oxygen consumption rates using the Oroboros O2k Oxygraph. (A) Time plot showing changes in absolute O_2_ concentration (left axis) and oxygen consumption rate (OCR, right axis) for the two conditions, following manual injections of oligomycin (oligo, 2 μM), uncoupling agent (FCCP, 1 μM per injection) and Antimycin A (AA, 2 μM). Glucose was used as the metabolic substrate at a concentration of 11.1 mM. (B) Comparison of Basal respiration, ATP-linked OCR, and Maximal (Max) respiration. Data are shown as mean ± SEM of n=4 biological replicates acquired in a single experiment and analyzed by unpaired t-test. Results are representative of n=12 biological replicates in total, from 4 independent experiments.

### ALA and DHA rewire monocyte metabolism towards glycolysis-derived ATP production

We then compared the effects of treating cells for 48h with Vehicle, ALA, DHA, or OA control (all at 40 μM) in a Seahorse XF mitochondrial stress test. Contrary to the previously reported finding where only ALA significantly altered monocyte bioenergetics in a clinical trial (Pauls et al., 2021), we found that both ALA and DHA reduced OCR parameters (basal, ATP-linked, maximal, and spare respiratory capacity, SRC) in THP-1 monocytes treated directly with these n-3 PUFA (**Figure 3A-B**). We also repeated the experiment with a more selective mitochondrial membrane uncoupler, BAM15 (**Supplementary Figure S1A)** and with a shorter, 24h treatment duration (**Supplementary Figure S1B)**, with consistent effects of ALA on OCR parameters. Interestingly, ECAR was unaffected by only 24 h treatment with ALA.

**Figure 3.**
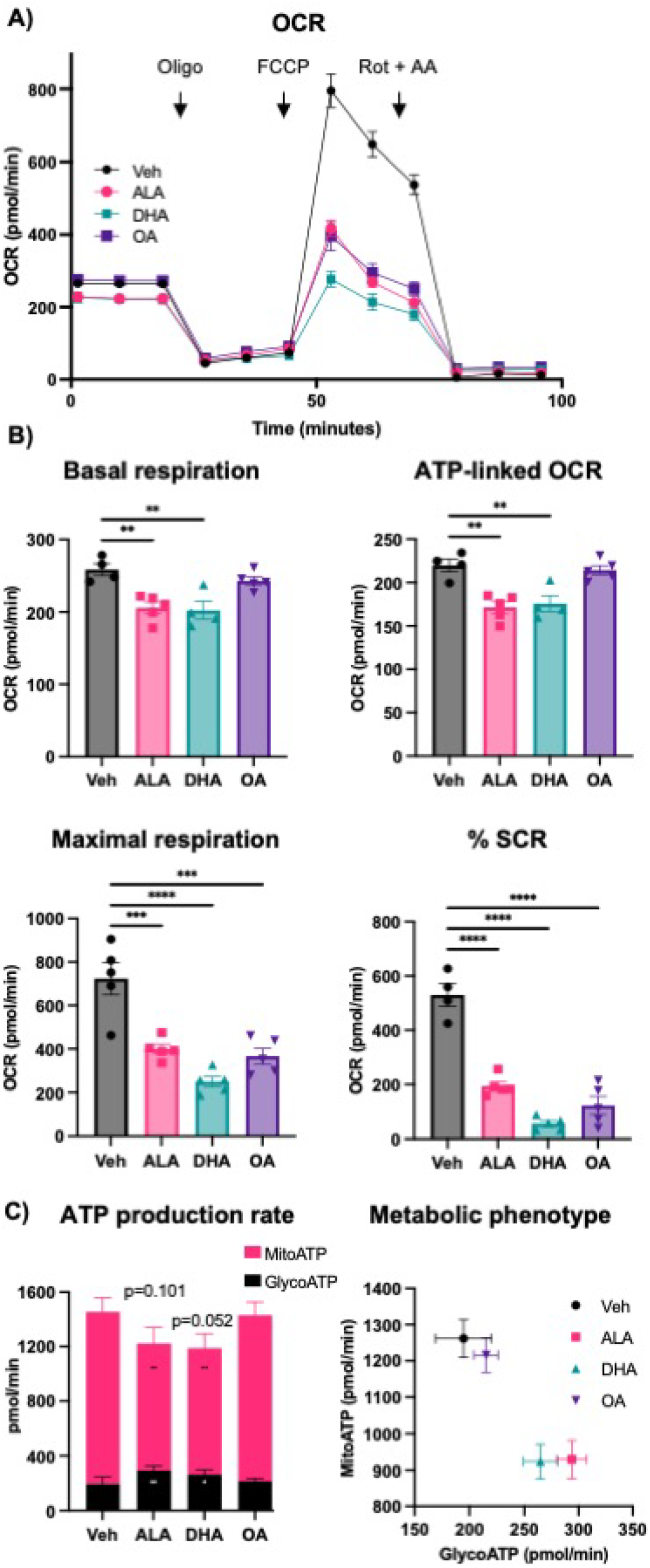
ALA and DHA induce a rewiring of monocyte ATP production. THP-1 cells were treated with vehicle or 40 μM of fatty acid (ALA, DHA, OA) for 48 hours with a bolus dose at 24 hours. A Seahorse XFe24 analyzer was used to assess bioenergetics. A) Time plot of a Mito Stress Test showing changes in Oxygen consumption rate (OCR) following the injection of Oligomycin (oligo, 1 μM), uncoupling agent (FCCP, 1.5 μM) and Rotenone + Antimycin A (Rot, 0.1 μM + AA, 1 μM). Glucose was used as the metabolic substrate at a concentration of 10 mM. B) Comparison of Basal respiration, ATP-linked OCR, Maximal respiration and Spare respiratory capacity (SRC) between treatments. C) Seahorse XF ATP rate Assays were performed to assess relative rates of ATP production via glycolytic (GlycoATP) and mitochondrial (MitoATP) pathways. Data are shown as mean ± SEM of n=4-6 technical replicate wells and analyzed via One-Way ANOVA; * P < 0.05, ** P < 0.01, *** P < 0.001, **** P < 0.0001. Results are representative of three independent experiments for each assay.

To better understand the overall energy phenotype of the cells, we further performed Seahorse ATP Rate assays. We saw that oxidative phosphorylation was responsible for the majority of ATP generation in these cells in all conditions (**Figure 3C**). Treatment with ALA and DHA, but not OA control reduced mitochondrial-derived ATP (MitoATP) and increased ATP generated from glycolysis (GlycoATP). Overall ATP generation trended downward but did not reach significance with either n-3 PUFA treatment. Vehicle and OA controls were tightly grouped in terms of metabolic phenotype, while ALA and DHA resulted in similar shifts in energy-generating pathways; ie. a lower reliance on oxidative phosphorylation and higher reliance on glycolysis for ATP generation, compared to the control conditions.

### ALA and DHA treatment lead to upregulation of pyruvate dehydrogenase kinase 4

Next, we investigated the effect of n-3 PUFA treatment on transcript levels of enzymes involved in regulating glucose catabolism. Our primary candidate was pyruvate dehydrogenase kinase 4 (*PDK4*), as it was reported to be upregulated in THP-1 cells by DHA according to a transcriptome microarray screen (Rodway et al., 2023). PDK4 phosphorylates and inhibits the activity pyruvate dehydrogenase (PDH), and therefore it is expected to negatively regulate oxidative phosphorylation (Jha et al., 2015; Zhang et al., 2014). Following a 48 h treatment with vehicle, ALA, DHA or OA, RNA was extracted and transcripts of interest were quantified by RT-qPCR. Both ALA and DHA treated cells had significantly higher *pdk4* mRNA levels compared to vehicle (**Figure 4**), although the increase was of much greater magnitude for DHA. Additionally, transcript levels of mitochondrial pyruvate carrier 1 (MPC1) and PDHA1 (the catalytic subunit of PDH) were investigated due to their roles in pyruvate transport and incorporation into the Kreb’s cycle, respectively. Neither candidate was significantly altered by the n-3 PUFA treatments.

**Figure 4.**
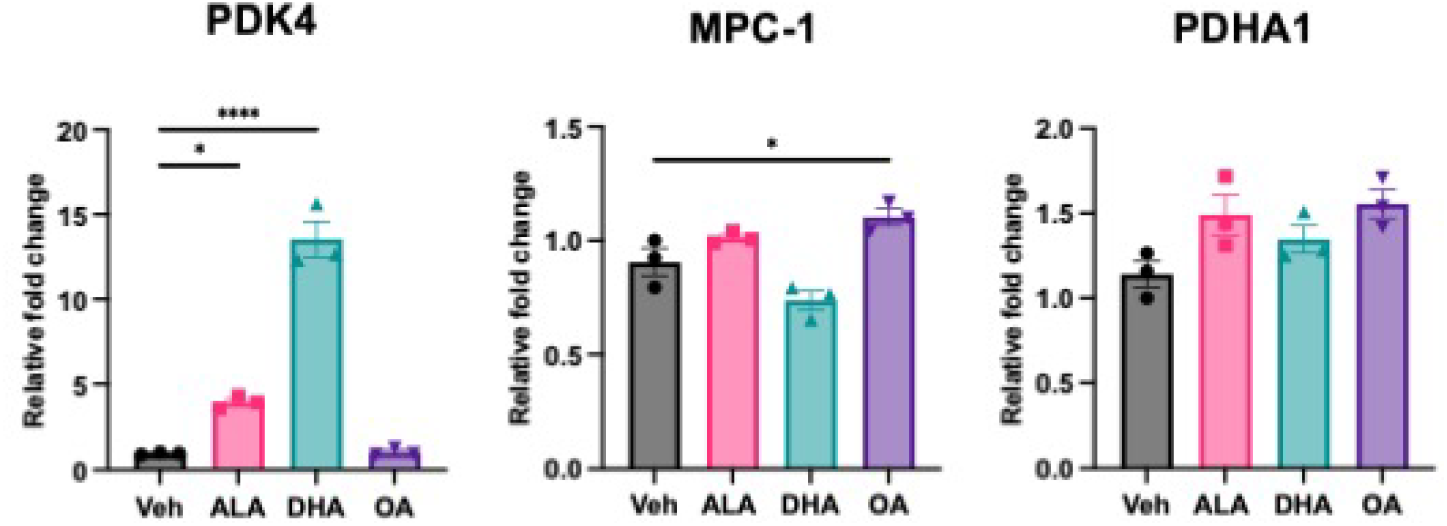
ALA and DHA increase PDK4 mRNA levels. THP-1 monocytes were treated with vehicle or 40 μM of fatty acid (ALA, DHA, OA) for 48 hours with a bolus at 24 hours. PDK4, MPC-1, and PDHA1 mRNA levels were assessed by RT-qPCR using the 2^-ddCt^ method with β-actin as the reference gene. Data are shown as mean ± SEM of three biological replicates, each the average of three technical replicate wells. Analysis is by One-Way ANOVA;* P < 0.05, **** P < 0.0001. Results are representative of three independent experiments for each transcript.

### DHA but not ALA increases reactive oxygen species

In non-hematopoietic cells, ALA and DHA have been reported to exhibit protective effects such as reducing reactive oxygen species (ROS) levels (Zhu et al., 2020; Sakai et al., 2017). In other studies, DHA increased mitochondrial ROS production (Shin et al., 2013). To investigate the effect of n-3 PUFA in our cellular model, we measured intracellular ROS production *via* flow cytometry using fluorescent indicators. After treating monocytes with 40 μM of fatty acids (ALA, DHA, OA) or vehicle, we used indicators for either total cellular ROS (CellROX) or mitochondrial superoxide (MitoSOX). We found that DHA-but not ALA-treated THP-1 monocytes had a significant increase in both cellular and mitochondrial ROS levels compared to vehicle and OA controls (**Figure 5**).

**Figure 5.**
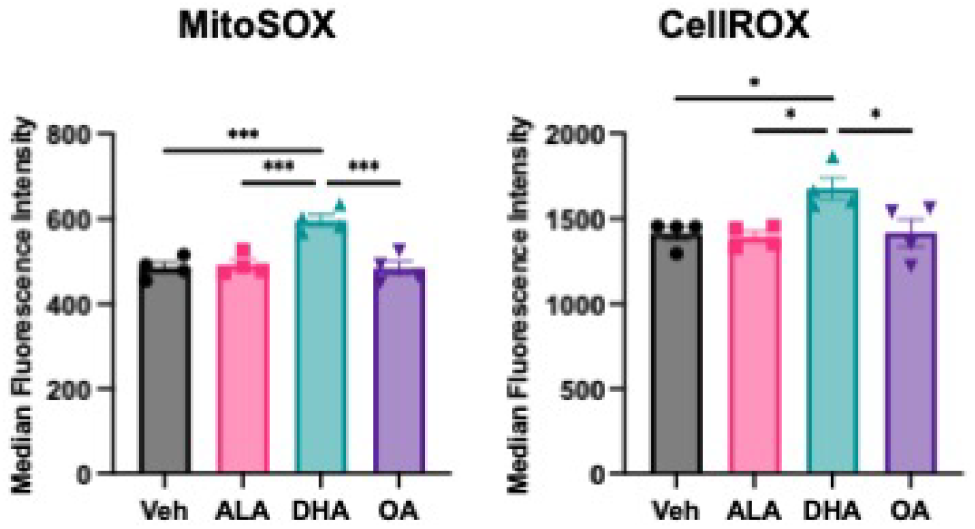
Treatment with DHA but not ALA increases reactive oxygen species. THP-1 monocytes were treated with either vehicle or 40 μM of fatty acid (ALA, DHA, OA) for 48 hours with a bolus dose at 24 hours. Cells were then stained with fluorescent indicator dyes for mitochondria-derived superoxide (MitoSOX, left), or total cellular reactive oxygen species (CellROX, right), quantified by flow cytometry. Results shown as median ± SEM of 4 biological replicates, analyzed via One-Way ANOVA: * P < 0.05, *** P < 0.001, **** P < 0.0001. Results are representative of three independent experiments for each indicator.

## Discussion

n-3 PUFAs have garnered much interest as a dietary or nutraceutical therapeutic that may reduce risk factors for metabolic diseases such as obesity and CVD (Faintuch et al., 2007; Itariu et al., 2012; Natto et al., 2019). This is in part due to its anti-inflammatory properties that have been demonstrated in a variety of pathologies and cell types. However, the mechanisms that dictate these beneficial effects are complex have yet to be fully elucidated. In this study, we identified a direct effect of n-3 PUFAs on glucose metabolism in THP-1 monocytes. Using both the Seahorse XF and Oroboros Oxygraph platforms, we demonstrate that ALA treatment (reaching significance at 40 μM) decreases basal, ATP-linked, and maximal respiration when glucose is provided as the metabolic substrate. Compared to Vehicle and OA control treatments, both ALA and DHA treatment reduced ATP production by oxidative phosphorylation and enhance ATP production by glycolysis to a similar extent. The overall impact may be a generally more “quiescent” metabolic state, although the observed qualitative reduction in total ATP generation did not quite reach the 95% confidence threshold. Furthermore, both n-3 PUFAs upregulated *pdk4* mRNA, but the magnitude of upregulation was much greater with DHA. Since PDK4 inhibits the PDH-mediated conversion of pyruvate to Acetyl-CoA, its activity is expected to both inhibit oxidative phosphorylation (as a result of less Acetyl Co-A entering the Kreb’s cycle) and promote glycolysis (as pyruvate is diverted to lactic acid, regenerating the electron carrier required for glycolysis). Thus, upregulation of *pdk4* may explain the metabolic rewiring effects of n-3 PUFA, or at least of DHA. Notably, while both n-3 PUFA had similar effects on bioenergetics, in the case of DHA, this was accompanied by an increase in mitochondrial and total ROS.

These results add to the complex, controversial and evolving landscape of innate immune cell immunometabolism research. Consistent across multiple innate immune cell types including monocytes, the pro-inflammatory TLR4 ligand LPS promotes glycolysis at the expense of oxidative phosphorylation (Krawczyk et al., 2010; Ó Maoldomhnaigh et al., 2021; Pauls et al., 2021). Thus, the idea that glycolysis promotes inflammatory activation was initially established. However, some recent studies in macrophages suggest that this may be entirely context-dependent. For example, the glycolytic biproduct lactic acid was demonstrated to support pro-resolving macrophage functions under some conditions (Schilperoort et al., 2023).

Relatively few studies have been performed specifically in monocytes that would help interpret the findings shown here. The study that inspired our analysis reported that, among women qualifying as obese (ie. BMI >30 and waist circumference over ethnicity-specific thresholds), higher BMI correlated to higher ATP-linked OCR and lower ECAR (Pauls et al., 2021). This was in line with an earlier clinical study that reported higher oxidative phosphorylation rates in peripheral blood mononuclear cells (PBMC, a mixed leukocyte population) from T2D participants compared to controls (Hartman et al., 2014). Along a similar line, a time-restricted feeding study demonstrated that fasting reduces both oxidative phosphorylation rates and inflammatory gene expression in mouse bone marrow monocytes, which also resist egress into circulation (Jordan et al., 2019).

This group of studies seem to contradict recent findings reported in a mouse model of obesity (Boroumand et al., 2022). These authors reported that high fat diet (HFD) feeding resulted in reduced oxidative phosphorylation in both Ly6C^low^ and Ly6C^high^ monocytes, as assessed by area-under-the-curve (AOC) measurement of the Mito Stress Test plot. HFD feeding also reduced glycolysis in Ly6C^low^ monocytes but increased glycolysis in Ly6C^high^ monocytes, as quantified by ECAR AOC. Although pro-inflammatory cytokines produced by these monocyte subsets were unchanged by HFD feeding, they found that white adipocyte conditioned medium promoted conversion of Ly6C^low^ to Ly6C^high^ cells, which are traditionally considered to be the more pro-inflammatory, infiltrating monocytes (Boroumand et al., 2022). Two additional studies suggest that oxidative phosphorylation, or at least PDH complex activity, may oppose inflammation. One group reported that PDK inhibition with dichloroacetate led to decreased protein levels of the pro-inflammatory cytokines IL-1β and TNF-α, as well as the anti-inflammatory cytokine IL-10 (Xuewei Zhu et al., 2020). Another group reported that mRNA levels of PDK4 were higher in CD14^+^ monocytes from coronary artery disease patients relative to healthy controls, and were further increased by exposure to low density lipoprotein (Du et al., 2021). Thus, there are multiple examples from literature where enhanced oxidative phosphorylation is speculated to be both anti-inflammatory and pro-inflammatory, depending on the context and, likely, the magnitude of change.

In the current study, the n-3 PUFA-induced shift away from oxidative phosphorylation towards glycolysis is not a complete reversal; oxidative phosphorylation remains the dominant pathway by which ATP is generated in these cells. Of note also is that ALA and DHA have consistently been described to have overall anti-inflammatory effects on immune cells (Hung et al., 2023; Pauls et al., 2018; Zhao et al., 2005). Now that a potential mechanism has been identified, work is ongoing to determine whether metabolic rewiring *via* PDK4 is responsible for any of the anti-inflammatory effects of ALA and DHA in monocytes.

In summary, our data suggest that the n-3 PUFAs ALA and DHA trigger a re-wiring of bioenergetic pathways in monocytes, promoting glycolysis at the expense of oxidative phosphorylation. This is possibly mediated *via* the upregulation of PKD4. Given the close relationship between cell metabolism and immune cell activation, this may represent a novel mechanism by which n-3 fatty acids modulate immune function and inflammation.

## Abbreviations

ALA: *α*-linolenic acid
BSA: Bovine serum albumin
CVD: Cardiovascular disease
DHA: Docosahexaenoic acid
ECAR: Extracellular acidification rate
ELISA: Enzyme-linked immunosorbent assay
ETC: Electron transport chain
LPS: lipopolysaccharide
MPC: Mitochondrial pyruvate carrier
OA: Oleic acid
OCR: oxygen consumption rate
PDH: Pyruvate dehydrogenase
PDK: Pyruvate dehydrogenase kinase
PUFA: polyunsaturated fatty acid
ROS: Reactive Oxygen Species
T2DM: Type 2 diabetes mellitus.

## Acknowledgements

We are extremely grateful to Dr. Versha Banerji and Maxim Skorodinsky of the University of Manitoba and the Paul Albrechtsen Research Institute - Cancer Care Manitoba for technical training and assistance with the Oroboros O2k Oxygraph measurements.

