## Supplementary Figure S1. for "Omega-3 polyunsaturated fatty acids modify glucose metabolism in THP-1 monocytes"

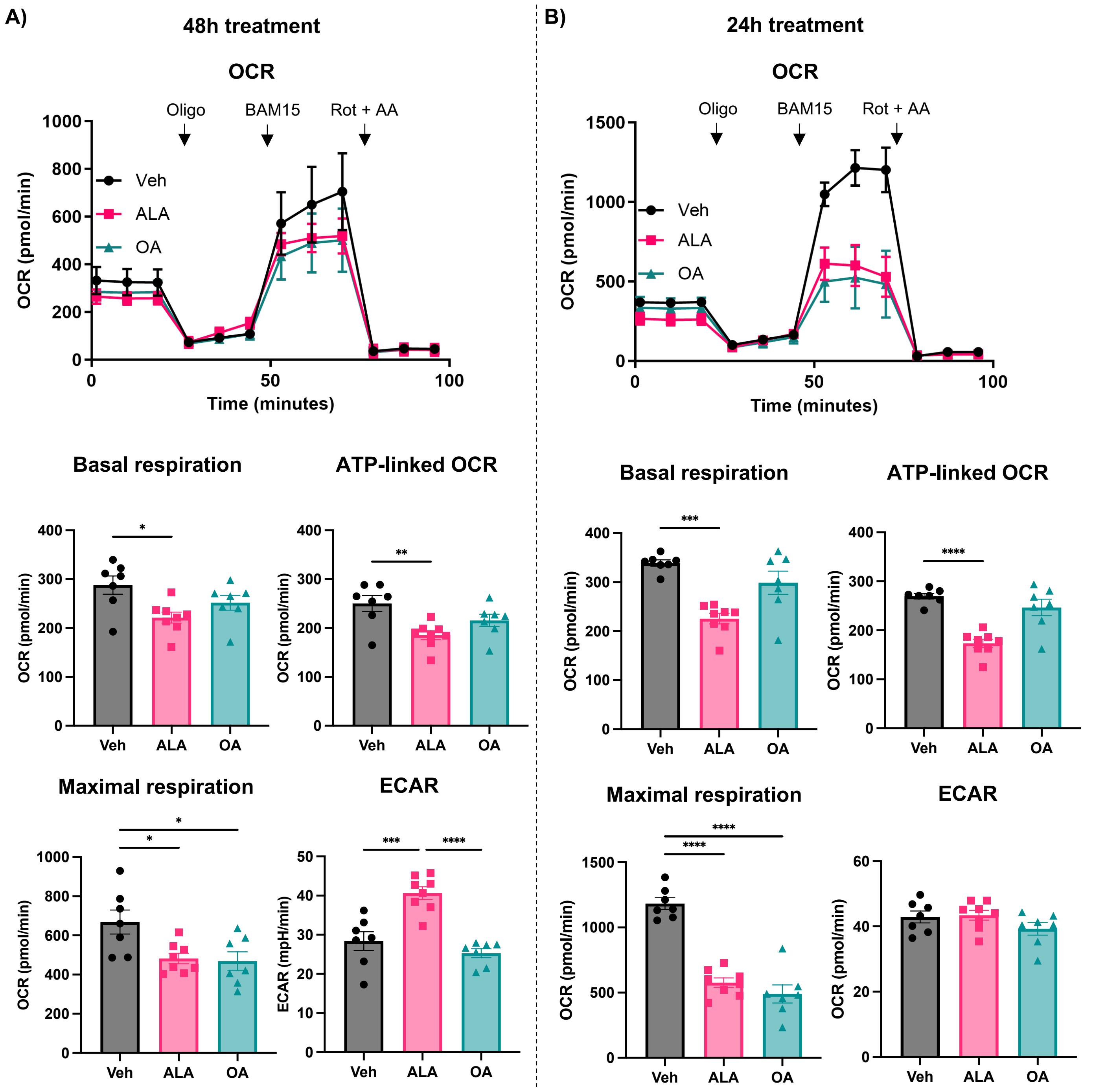


**Supplementary Figure S1. ALA alters bioenergetic parameters after only 24h, including Maximal respiration assessed using BAM15 as the uncoupling agent.** THP-1 cells were treated with vehicle or 40 µM of fatty acid (ALA or OA) for 48 hours with a bolus dose at 24 hours (A) or for 24h (B). A Seahorse XFe24 analyzer was used to assess bioenergetics. A) Time plot of a Mito Stress Test showing changes in Oxygen consumption rate (OCR) following the injection of Oligomycin (oligo, 1 µM), uncoupling agent (BAM15, 1.5 µM) and Rotenone + Antimycin A (Rot, 0.1 µM + AA, 1 µM). Glucose was used as the metabolic substrate at a concentration of 10 mM. Comparisons between groups are shown for Basal respiration, ATP-linked OCR, Maximal respiration and extracellular acidification rate (ECAR). Data are shown as mean ± SEM of n=4-6 technical replicate wells and analyzed via One-Way ANOVA; * $P < 0.05$, ** $P < 0.01$, *** $P < 0.001$, **** $P < 0.0001$.
